# USP29 is a novel non-canonical Hypoxia Inducible Factor-α activator

**DOI:** 10.1101/2020.02.20.957688

**Authors:** Amelie S Schober, Inés Martín-Barros, Teresa Martín-Mateos, Encarnación Pérez-Andrés, Onintza Carlevaris, Sara Pozo, Ana R Cortazar, Ana M Aransay, Arkaitz Carracedo, Ugo Mayor, Violaine Sée, Edurne Berra

## Abstract

Hypoxia Inducible Factor (HIF) is the master transcriptional regulator that orchestrates cellular adaptation to low oxygen. HIF is tightly regulated via the stability of its α-subunit, which is subjected to oxygen-dependent proline hydroxylation by Prolyl-Hydroxylase Domain containing proteins (PHDs/EGLNs), and ultimately targeted for proteasomal degradation through poly-ubiquitination by von-Hippel-Lindau protein (pVHL). However, sustained HIF-α signalling is found in many tumours independently of oxygen availability pointing towards the relevance of non-canonical HIF-α regulators. In this study, we establish the Ubiquitin Specific Protease 29 (USP29) as direct post-translational activator of HIF-α in a variety of cancer cell lines. USP29 binds to HIF-α, decreases poly-ubiquitination and thus protects HIF-α from proteasomal degradation. Deubiquitinating activity of USP29 is essential to stabilise not only HIF-1α but also HIF-2α, via their C-termini in an oxygen/PHD/pVHL-independent manner. Furthermore, in prostate cancer samples the expression of *USP29* correlates with the HIF-target gene *CA9* (carbonic anhydrase 9) as well as disease progression and severity.

## Introduction

Besides being an essential developmental and physiological stimulus, hypoxia is associated with pathologies such as cancer, metabolic, inflammatory, neurodegenerative and ischemic diseases. Hypoxia is indeed a feature of most human cancers (Semenza, 2012). Hence, cancer cells and their environment need to adapt to and survive under low oxygen availability.

The transcription factor HIF (hypoxia-inducible factor) is the central regulator of the adaptive cellular program in response to limited oxygen availability. The two HIF subunits, HIF-α and HIF-β are constitutively expressed, but the stability of HIF-α protein is tightly regulated through the ubiquitin-proteasome system (UPS) in order to avoid inadequate HIF signalling (Huang et al, 1996). In well-oxygenated cells, HIF-α is hydroxylated by the oxygen sensors PHDs/EGLNs, and subsequently ubiquitinated by the ubiquitin E3-ligase von-Hippel-Lindau protein (pVHL) (Bruick & McKnight, 2001; Epstein et al, 2001; Ivan et al, 2001; Jaakkola et al,2001; Maxwell et al, 1999). Ubiquitinated HIF-α protein is degraded by the proteasome (Salceda & Caro, 1997). Upon hypoxia, PHDs/EGLNs activity is compromised, HIF-α escapes from degradation, dimerises with HIF-β, binds to RCGTG motives (hypoxia responsive elements, HRE) within the regulatory domains of target genes and transcriptionally drives their expression (Arany et al, 1996; Wang & Semenza, 1993). HIF-targets involve among many others, genes that enhance glycolysis and metabolic rewiring, angiogenesis and resistance to apoptosis (Schodel et al, 2011). Accordingly, sustained expression of HIF-α in tumours has been associated with higher aggressiveness, migratory and metastasis-initiating potential and therefore worse prognosis (Trastour et al, 2007; Zhong et al, 1999). However, HIF-α stabilisation does not always correlate with tissue oxygenation (Mayer et al, 2008).

Especially in the context of cancer, additional UPS related proteins have been described to be involved in the control of HIF-α stability (recently reviewed in (Schober & Berra, 2016)). Among those are the HIF-α destabilisers RACK1, MDM2, Fbw7 and CHIP that control HIF-α stability in a non-canonical way, namely independently of O_2_/PHDs and/or pVHL. Of the family of deubiquitinating enzymes (DUBs), able to specifically deconjugate ubiquitin from targeted proteins, USP20 (also called pVHL interacting deubiquitinating enzyme 2, VDU2), MCPIP1, USP8 and UCHL1 emerged as new HIF-α regulators, as they reverse the canonical HIF-αubiquitination. Furthermore, USP28 antagonizes Fbw7-mediated HIF-1α degradation, and Cezanne (OTUD7B) protects HIF-1α from lysosomal degradation, and are therefore implicated in the non-canonical HIF-1α regulation (Altun et al, 2012; Bremm et al, 2014; Flugel et al, 2012; Goto et al, 2015; Li et al, 2005; Roy et al, 2013; Troilo et al, 2014). Surprisingly, to date no DUB has been shown to exhibit hydrolase activity towards HIF-2α.

In 2000 a new gene in the Peg3 (paternally expressed gene 3) region was discovered. Like all genes in this region, it was shown to be imprinted and, in this specific case, to be paternally expressed. Due to its structural homology with the ubiquitin specific proteases (USPs), the biggest class of DUBs, it was named USP29. USP29 mRNA was only detectable by Northern Blot in murine brain and in testis of mice and humans (Kim et al, 2000). It was not until 2011 that the first biological function of the 922 aa long USP29 gene product was described and showed that H_2_O_2_ treatment induced the expression of USP29 (Liu et al, 2011). They reported that USP29 bound to p53 and stabilised it by decreasing its ubiquitination. A few years later, USP29 was also described to bind to the cell cycle checkpoint adapter claspin and that USP29 silencing reduced basal claspin levels (Martin et al, 2015). Here, we show that USP29 is a novel non-canonical regulator of both HIF-1α and HIF-2α. USP29 binds to HIF-α in an oxygen-independent manner, deubiquitinates it and therefore rescues HIF-α from proteasomal degradation.

## Results

### USP29 is a positive regulator of HIF-1α

The hypoxia pathway is under exquisite control by reversible ubiquitination. In order to identify hypoxia specific deubiquitinating enzymes (DUBs), we carried out an unbiased loss-of function screen using pools of small hairpin RNAs (shRNAs) to individually inhibit the expression of 66 human DUBs and the hypoxia-driven LUC reporter. USP29 came up as one of the strongest hits from three independent screenings carried out in triplicates. Indeed, the silencing of endogenous *USP29* with a pool of 3 independent shRNAs in HeLa cells significantly reduced the hypoxia-driven HRE-luciferase expression (Figure 1A). In concordance with this data, silencing of endogenous *USP29* also abrogated the hypoxic induction of the HIF target gene *CA9* (Figure 1B). The pool of shUSP29s efficiently silenced GFP-USP29 at mRNA and protein levels (Expanded View Figure 1A). Interestingly, in cells silenced for endogenous *USP29* the accumulation of HIF-1α protein in hypoxia was significantly decreased and the induction of CAIX and PHD2 was impaired to a similar extent as when silencing *HIF1A* (Figure 1C). Anyhow, *HIF1A* mRNA was not affected by the silencing of *USP29* (Expanded View Figure 1B). More importantly, similar to the pan-hydroxylase inhibitor DMOG, the ectopic expression of USP29 led to the accumulation of endogenous HIF-1α, CAIX and PHD2 even in normoxia (Figure 1D). Nonetheless, *HIF1A* mRNA expression was not affected by the USP29 overexpression (Expanded View Figure 1C), pointing to USP29 as a novel upstream post-translational activator of HIF-1α.

**Figure 1.**
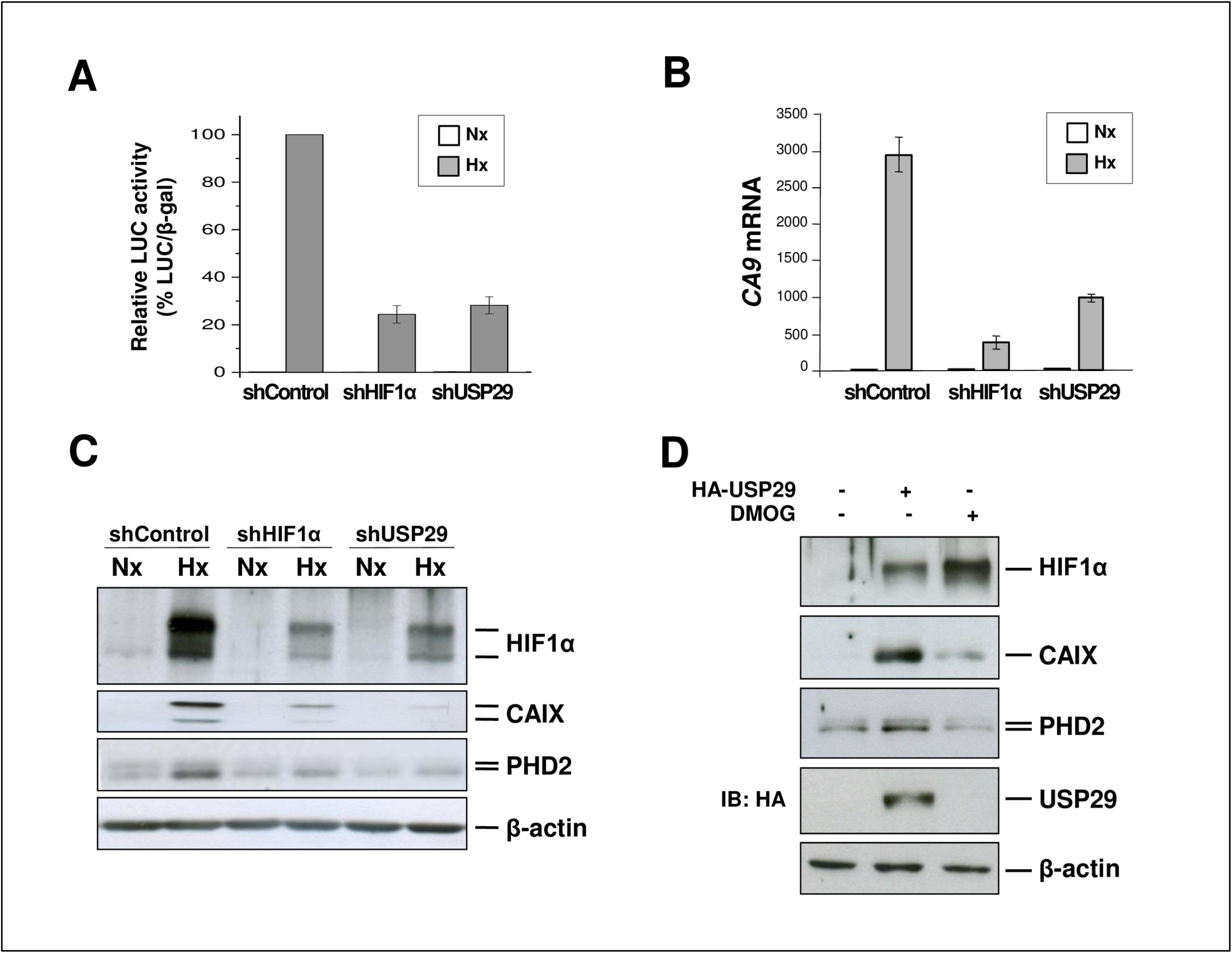
USP29 is a positive regulator of HIF-1α. **A** HeLa cells were silenced with scrambled or shRNAs targeting *HIF1A* and *USP29* and transfected with a reporter vector (pRE-Δtk-Luc) containing three copies of the HRE from the erythropoietin gene and CMV-β-gal to normalize for transfection efficiency. Cells were incubated for 16 h in normoxia (21% O_2_) or hypoxia (1% O_2_) and luciferase and β-galactosidase activities were measured. **B** HeLa cells were treated as in A and total RNA was extracted, reverse-transcribed and expression of *CA9* was determined by qPCR. **C** Whole cell extracts (WCE) from HeLa cells treated as in A were subjected to SDS-PAGE followed by immunoblotting with the indicated antibodies. **D** HEK293T cells were transfected with empty vector or HA-USP29 and left untreated or treated with DMOG for 4 hours prior to lysis. WCE were subjected to SDS-PAGE and immunoblotting was performed using the indicated antibodies.

### USP29 upregulates HIF-1α in a non-canonical way

Surprisingly, the HIF-1α that accumulated in the presence of USP29 in normoxic conditions induced PHD2 and CAIX (Figure 1D), albeit being prolyl-hydroxylated (Figure 2A). Furthermore, the ectopic expression of USP29 also accumulated HIF-1α DM^(PP/AA)^, a HIF-1α mutant whose two oxygen-sensitive proline residues have been replaced by alanines (P402/564A), suggesting that USP29 regulates HIF-1α in a non-canonical way (Figure 2B).

**Figure 2.**
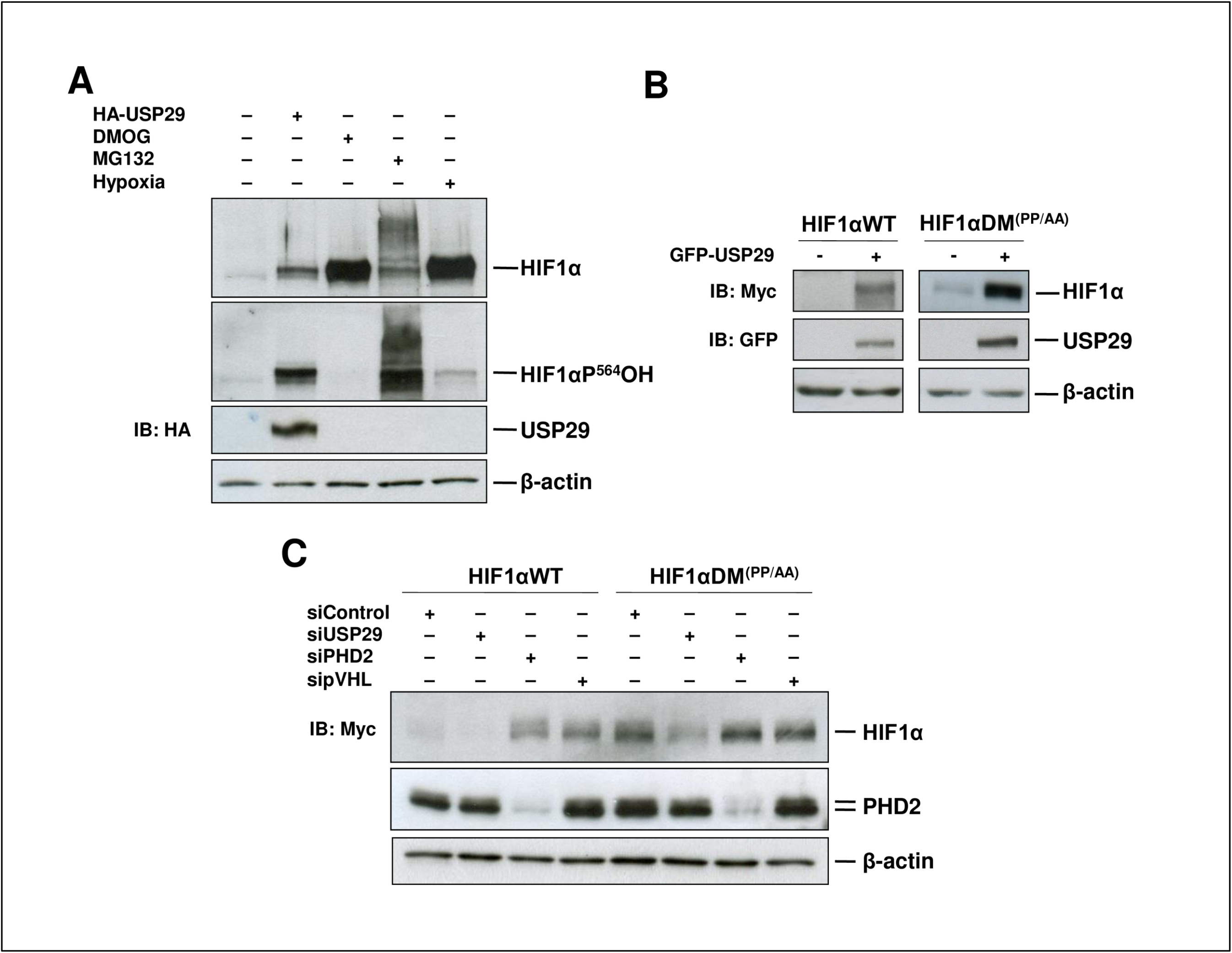
USP29 regulates HIF-1α in a non-canonical way. **A** HEK293T cells were transfected with empty vector or HA-USP29 and treated with the hypoxia mimetic DMOG (1 mM), the proteasome inhibitor MG132 (10 μM) or hypoxia (1% O_2_) for 4 hours. WCE were prepared and analysed by immunoblotting with the indicated antibodies. **B** HEK293T cells were co-transfected with myc-HIF-1α or myc-HIF-1α DM^(PP/AA)^ and empty vector or GFP-USP29. Levels of myc- and GFP-tagged proteins in WCE were determined by immunoblotting in WCE. **C** HEK293T cells were silenced with control or siRNAs (20 nM) targeting endogenous *USP29*, *PHD2/EGLN1* or *pVHL* mRNA and transfected with myc-HIF-1α or myc-HIF-1α DM^(PP/AA)^. Total cell extracts were subjected to SDS-PAGE followed by immunoblotting with the indicated antibodies.

Consistently, silencing of endogenous *USP29* with 2 different siRNA sequences decreased both, HIF-1α WT and HIF-1α DM^(PP/AA)^ protein levels (Figure 2C and Expanded View Figure 2A). As expected, the silencing of the canonical negative regulators, *PHD2/EGLN1* and *pVHL*, only affected HIF-1α WT but not HIF-1α DM^(PP/AA)^ (Figure 2C). Similarly, the overexpression of the Ub E3-ligase pVHL did only affect HIF-1α WT, but not the DM protein (Expanded View Figure 2B). Taken together, these results indicate that USP29 acts on HIF-1α through a non-canonical mechanism.

### Universality of USP29’s effect on HIF-α

The effect of USP29 on HIF-1α was observed in a variety of cell lines of different origins, including A2780 (ovarian cancer), PC3 and LnCaP (prostate cancer), SH-SY5Y and SK-N-AS (neuroblastoma) and MDA-MB-231 (breast cancer). In all tested cell lines the overexpression of USP29 led to an increase in HIF-1α DM^(PP/AA)^ levels (Figure 3A), indicating that this regulation might be a wide phenomenon. Interestingly, not only HIF-1α but also both, the wild type and the oxygen-insensitive DM^(PP/AA)^ forms of HIF-2α/EPAS accumulated upon overexpression of USP29 (Figure 3B).

**Figure 3.**
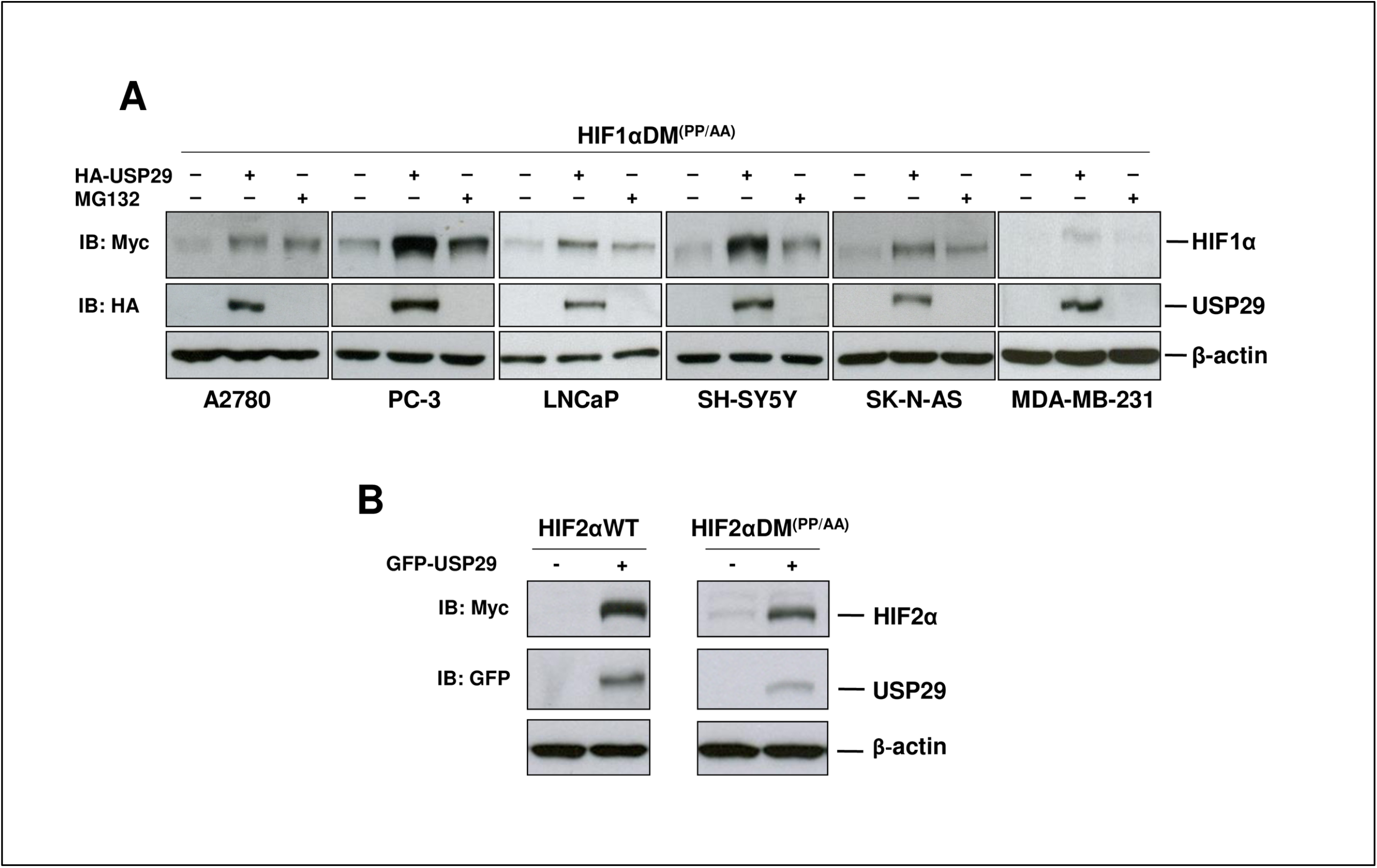
Wide impact of USP29 on HIF-α. **A** Cancer cell lines of different origins were co-transfected with myc-HIF-1α DM^(PP/AA)^ and empty vector or HA-USP29 and left untreated or treated with the proteasome inhibitor MG132 (10 μM) for 4 hours. WCE were subjected to SDS-PAGE followed by immunoblotting with the indicated antibodies. **B** HEK293T cells were co-transfected with myc-HIF-2α or myc-HIF-2α DM^(PP/AA)^ and empty vector or GFP-USP29 and total cell extracts were analysed by immunoblotting.

### USP29 stabilises HIF-α by protecting it from proteasomal degradation

In order to determine the molecular mechanism of HIF-α DM^(PP/AA)^ accumulation by USP29, we treated HEK293T cells with the proteasome inhibitor MG132 for 4 hours in the absence or presence of ectopic USP29 (Figure 4A). Both, the USP29 overexpression and the proteasome inhibition induced HIF-1α DM^(PP/AA)^ accumulation, but the lack of additivity indicated that they both acted on the same pathway. Furthermore, as USP29 accumulated HIF-1α DM^(PP/AA)^ more efficiently than MG132, we tested whether HIF-1α DM^(PP/AA)^ was also degraded via the lysosomal pathway. Yet, the inhibition of this pathway by treatment with chloroquine failed to prevent HIF-1α DM^(PP/AA)^ degradation (Expanded View Figure 3A), confirming that it requires the proteasome activity and suggesting that the difference between MG132- and USP29-induced HIF-1α DM^(PP/AA)^ accumulation was due to incomplete proteasome inhibition.

**Figure 4.**
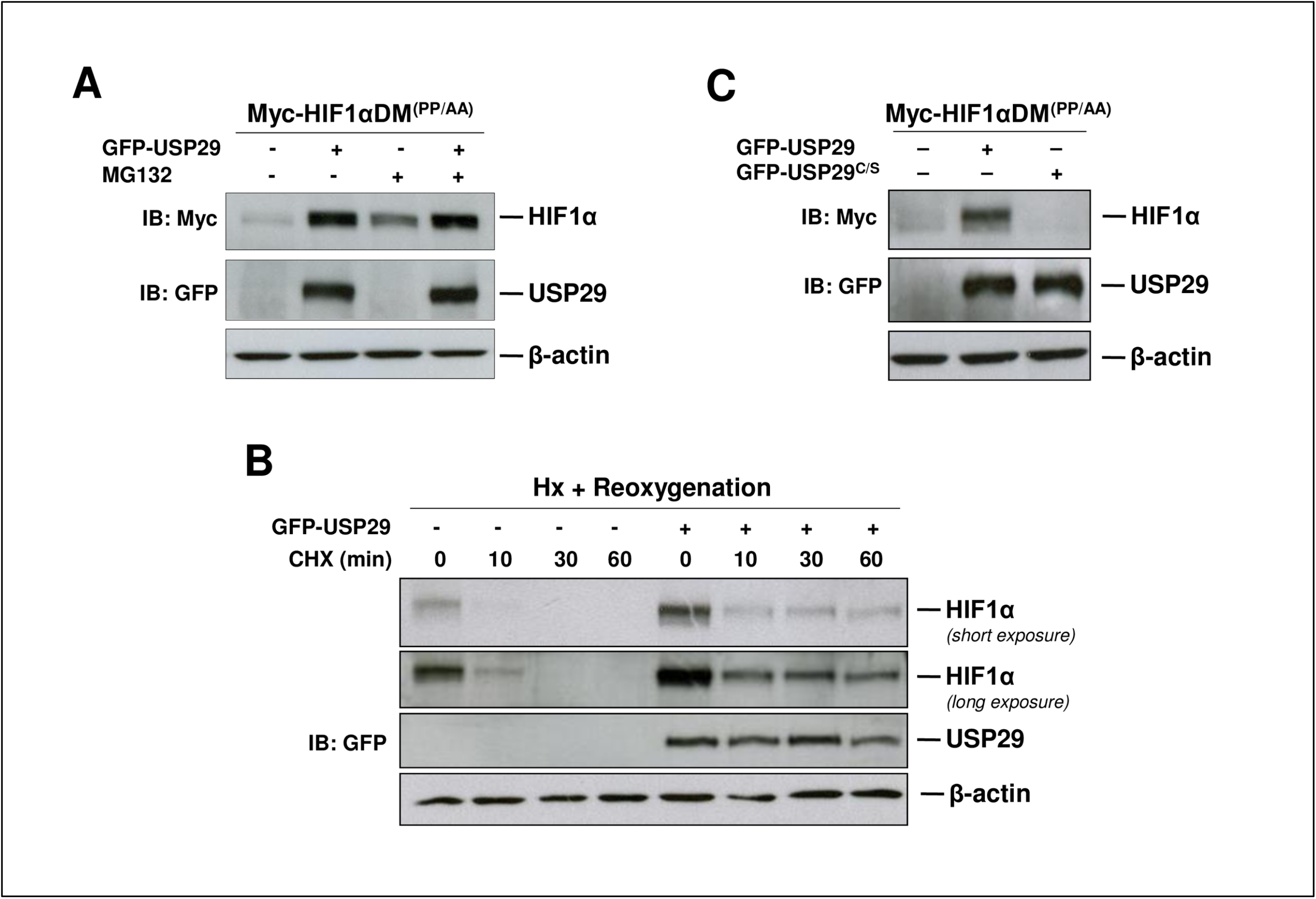
USP29 stabilises HIF-1α by protecting it from proteasome-mediated degradation. **A** HEK293T cells were co-transfected with myc-HIF-1α DM^(PP/AA)^ and empty vector or GFP-USP29 and left untreated or treated with the proteasome inhibitor MG132 (10 μM) for 4 hours. Total cell extracts were subjected to SDS-PAGE followed by immunoblotting with the indicated antibodies. **B** HEK293T cells were transfected with empty vector or GFP-USP29 and incubated in hypoxia (1% O_2_) for 4 hours. Then cells were treated with cycloheximide (20 μg/ml) to inhibit protein synthesis, reoxygenated and cell extracts were prepared at the indicated time points. HIF-1α protein levels were determined by Western Blotting. **C** HEK293T cells were co-transfected with myc-HIF-1α DM^(PP/AA)^ and empty vector, GFP-USP29 or GFP-USP29^C/S^. Cell extracts were subjected to immunoblotting with the indicated antibodies.

Cycloheximide experiments showed that USP29 increased HIF-1α DM^(PP/AA)^’s half-life from ≅1 to ≅3 hours (Expanded View Figure 3B). More importantly, USP29 stabilised endogenous HIF-1α upon reoxygenation (Figure 4B). Although USP29 did not avoid the initial HIF-1α degradation within the first 10 minutes of reoxygenation, thereafter HIF-1α levels remained stable during at least one hour in the presence of USP29, while the protein was not longer detectable 30 minutes after reoxygenation in the absence of USP29. To gain further insight into how USP29 stabilised HIF-α, we generated a catalytically inactive USP29 mutant by replacing its active site cysteine residue C294 with a serine (USP29^C/S^). This mutation completely abrogated USP29’s ability to accumulate HIF-1α DM^(PP/AA)^ (Figure 4C), pointing towards a crucial role of USP29’s ubiquitin specific peptidase activity in HIF-α DM^(PP/AA)^ stabilisation.

### USP29 interacts with and deubiquitinates HIF-α DM^(PP/AA)^

As the catalytical activity of USPs is responsible for removal of (poly)ubiquitin chains from their target proteins, we next tested whether USP29 was able to function as a deubiquitinase for HIF-α. We first analysed USP29 and HIF-α interaction using fluorescence lifetime based FRET measurements. The fluorescence lifetime of the FRET donor, Clover-HIF-1α DM^(PP/AA)^, was significantly decreased from 2.86 ± 0.02 ns to 2.7 ± 0.09 ns in the presence of the FRET acceptor mCherry-USP29 (Figure 5A). As FRET only occurs when both fluorophores are in very close proximity (around 6 nm), these data clearly show that USP29 is directly bound to HIF-1α DM^(PP/AA)^. Similar results were obtained when we analysed the interaction between USP29 and HIF-2α DM^(PP/AA)^ (Expanded View Figure 4A). HIF-2α DM^(PP/AA)^-GFP’s lifetime was significantly reduced from 2.39 ± 0.01 ns to 2.28 ± 0.06 ns in the presence of the FRET acceptor mCherry-USP29. Furthermore, when GFP-tagged HIF-1α DM^(PP/AA)^ or GFP alone were immunoprecipitated from HEK293T cells, we found HA-USP29 to interact with GFP-tagged HIF-1α DM^(PP/AA)^, but not with GFP alone (Expanded View Figure 4B). Next, we cotransfected GFP-tagged HIF-1α DM^(PP/AA)^ together with FLAG-ubiquitin either in the absence or the presence of HA-USP29 or HA-USP29^C/S^. After the enrichment of the ubiquitinated proteome by MG132-treatment, GFP-HIF-1α DM^(PP/AA)^ was pulled-down under highly denaturing conditions and anti-FLAG-antibody was used to detect ubiquitinated GFP-HIF-1α DM^(PP/AA)^. We found that USP29 wild type, but not the catalytically inactive USP29^C/S^, considerably decreased the basal ubiquitination of HIF-1α DM^(PP/AA)^ and increased the non-modified population of HIF-1α DM^(PP/AA)^ (Figure 5B). Accordingly, when silencing endogenous *USP29*, we observed increased poly-ubiquitination of HIF-1α DM^(PP/AA)^ (Figure 5C), pointing towards a basal deubiquitinating activity of endogenous USP29. Expression of a siRNA-resistant USP29 restored the basal HIF-1α DM^(PP/AA)^ ubiquitination pattern (Figure 5C right lane). Furthermore and in concordance with Fig 3B, USP29 also exerted deubiquitination activity towards HIF-2α DM^(PP/AA)^ (Expanded View Figure 4B). Taken together, our results indicate that endogenous and ectopic USP29 is an efficient deubiquitinase for HIF-α DM^(PP/AA)^ thereby increasing HIF-α stabilisation and subsequent HIF activation.

**Figure 5.**
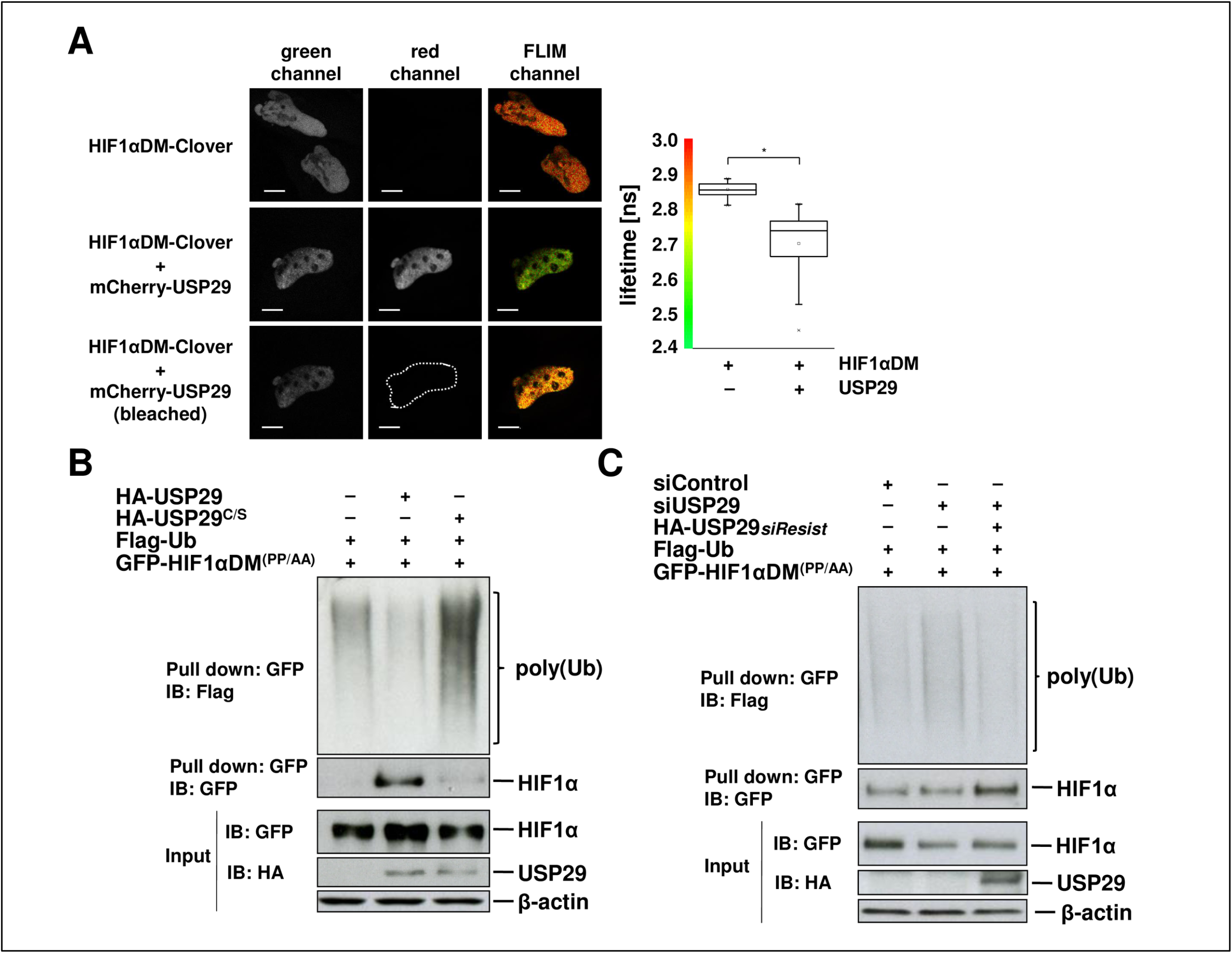
USP29 deubiquitinates HIF-α DM^(PP/AA)^. **A** HeLa cells were transfected with the FRET donor Clover-HIF-1α DM^(PP/AA)^ alone or together with the FRET acceptor mCherry-USP29. Fluorescence images for donor (green) and acceptor (red) channel were acquired (left and central panel). The lifetime of the donor was measured and pseudo-colour coded fluorescence life time images (FLIM) were generated. From 3 independent experiments average lifetimes of the donor in the absence (n = 34) and the presence (n = 25) of the FRET acceptor were calculated. Scale bars are 10 μm long, (*) p = 1.32*10^-8^. **B** HEK293T cells were co-transfected with GFP-HIF-1α DM^(PP/AA)^, FLAG-ubiquitin and either HA-USP29 or HA-USP29^C/S^. Cells were treated with the proteasome inhibitor MG132 (10 μM) for 2 hours and lysed in the presence of the DUB inhibitor NEM. GFP-HIF-1α DM^(PP/AA)^ was pulled down with GFP-traps^®^ and subjected to stringent washes (8 M urea, 1% SDS). Ubiquitinated and non-ubiquitinated GFP-HIF-1α DM^(PP/AA)^ protein in the eluate was analysed by immunoblotting with anti-FLAG and anti-GFP antibodies, respectively. **C** HEK293T cells were silenced with a control or a siRNA targeting *USP29* (20 nM) and co-transfected with GFP-HIF-1α DM^(PP/AA)^, FLAG-ubiquitin and either empty vector or siRNA-resistant HA-USP29. Treatment of cells, pull-down with GFP-traps^®^ and subsequent analysis of the ubiquitinated and non-ubiquitinated GFP-HIF-1α DM^(PP/AA)^ protein in the eluate were performed as in B.

### USP29 targets the C-terminal part of HIF-α

To identify the potential lysine residues targeted by USP29’s deubiquitinating activity, we tested several truncated forms of HIF-1α DM^(PP/AA)^ for their susceptibility to USP29. The N-terminal part, HIF1αDM^1-657^, was not affected by the presence of USP29, while the C-terminal end (HIF-1α^630-826^) accumulated in the presence of USP29 similarly to the full-length protein (Figure 6A). The USP29^C/S^ mutant that lacked catalytical activity was not able to accumulate HIF-1α^630-826^ (Expanded View Figure 5A). Correspondingly, USP29 acted also on the C-terminus of HIF-2α (Expanded View Figure 5B). We used truncations of the C-terminus to further confine the USP29 target site within HIF-1α. HIF-1α^630-713^ and HIF-1α^630-750^ were resistant to USP29-mediated accumulation (Expanded View Figure 5C) and pointed out the importance of the very C-terminal tail of HIF-1α for this regulation. This tail contains two evolutionary conserved lysines (K752 and K755), which are also shared by HIF-2α and a neighbouring lysine (K758) (Expanded View Figure 5D). Mutation of all three lysines to arginines (HIF-1α DM^KKK/RRR^) conferred to this mutated protein a higher stability in cycloheximide experiments (Figure 6B). Importantly, the basal ubiquitination of HIF-1α DM^KKK/RRR^ was significantly reduced as compared to HIF-1α DM (Figure 6C).

**Figure 6.**
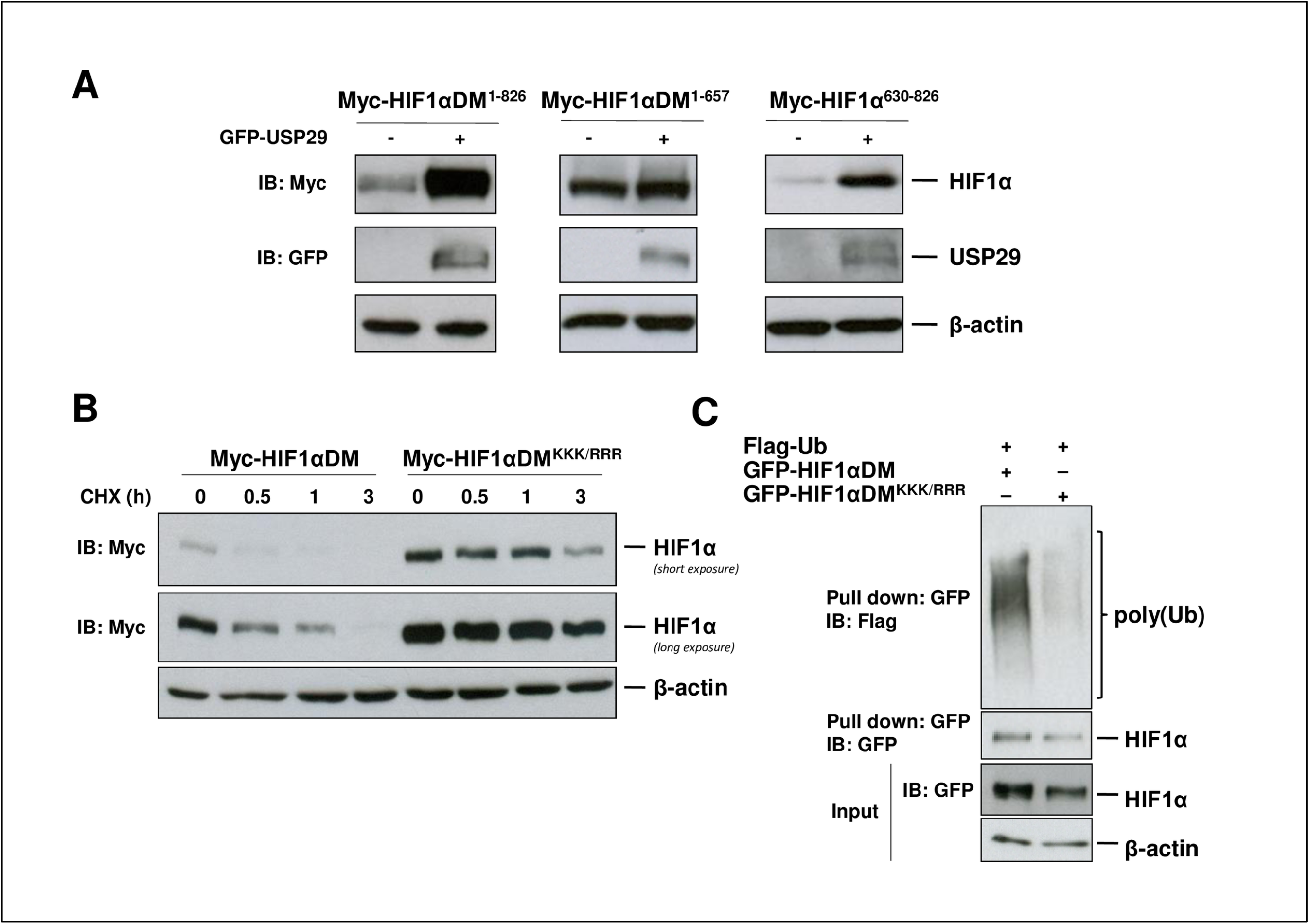
USP29 targets the C-terminal part of HIF-α. **A** HEK293T cells were co-transfected with myc-HIF-1α DM^1-826^, myc-HIF-1α DM^1-657^ or myc-HIF-1α DM^630-826^ and either empty vector or GFP-USP29. Whole cell extracts were subjected to SDS-PAGE followed by immunoblotting with the indicated antibodies. **B** HEK293T cells were transfected with myc-HIF-1α DM or myc-HIF-1α DM^KKK/RRR^ and cells were treated with cycloheximide (CHX) (20 μg/ml) to inhibit protein synthesis. Cell extracts were collected at the indicated times after CHX treatment and protein levels of the myc-tagged HIF-1α DM^(PP/AA)^ proteins were analysed by western blot. **C** HEK293T cells were co-transfected with GFP-HIF-1α DM or myc-HIF-1α DM^KKK/RRR^ and FLAG-ubiquitin. Cells were treated with the proteasome inhibitor MG132 (10 μM) for 2 hours and lysed in the presence of the DUB inhibitor NEM. GFP-tagged protein was pulled down with GFP-traps^®^ and subjected to stringent washes (8 M urea, 1% SDS). Ubiquitinated and non-ubiquitinated GFP-HIF-1α protein in the eluate was analysed by immunoblotting with anti-FLAG and anti-GFP antibodies, respectively.

### USP29 levels correlate with tumour progression and HIF target gene expression

The fact that USP29 stabilises HIF-α and is able to maintain hypoxia signalling switched on in normoxic conditions, led us to inquire its potential function in tumour progression. We therefore assessed whether USP29 expression was altered in certain tumours. Data mining analysis of publicly available databases revealed that *USP29* expression was significantly correlated with prostate cancer progression (Figure 7A). The expression levels of *USP29* mRNA increased from normal tissue over primary tumour to metastasis. Interestingly, *USP29* expression exhibited a significant association with the Gleason Score (GS), used in the clinics to stratify prostate cancer patients and predict their prognosis, as reflected by higher GS associated with higher *USP29* expression levels (Figure 7B). Furthermore, in the prostate cancer samples the expression of *USP29* also showed a significant positive correlation with the expression levels of the HIF target gene *CA9* (Figure 7C).

**Figure 7.**
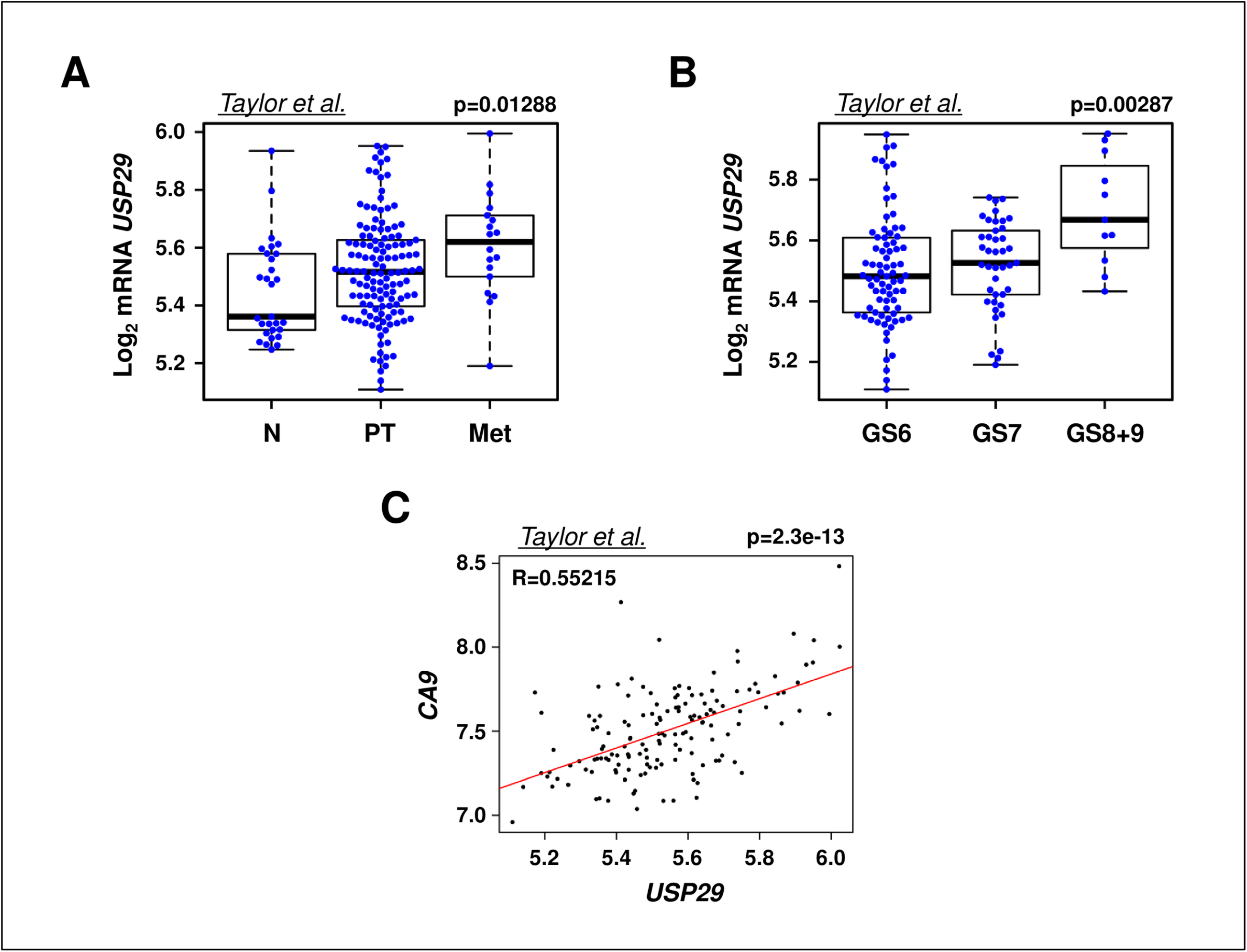
USP29 expression in prostate cancer. **A, B** Gene expression analysis of *USP29* in a dataset of prostate cancer samples (Taylor et al, 2010). *USP29* mRNA levels in prostate samples were compared on the basis of their tissue origin (A) or the Gleason score (GS) of the patient (B) (normal tissue (N): n = 29, primary tumours (PT): n = 131; metastatic tumours (Met): n = 19). **C** Correlation analysis between *USP29* and *CA9* mRNA levels in the aforementioned dataset of primary prostate cancer samples (Taylor, n = 131).

## Discussion

As the master transcription factor for hypoxia induced genes, HIF is the central component of cellular oxygen sensing. However, the pathway can be active even in the absence of hypoxia and HIF-α expression does not always correlate with tissue oxygenation. Notably, sustained HIF signalling occurs in many pathological conditions including cancer and inflammatory diseases pointing towards the relevance of non-canonical regulators of the HIF pathway. In the present study, we reported a novel insight into these regulatory mechanisms via USP29. We provided clear evidence that the ubiquitin specific protease 29 (USP29) is a new non-canonical and direct positive regulator of HIF-α stability in a panel of different cell lines. USP29 bound to poly-ubiquitinated HIF-α, is responsible for its deubiquitination and hence protects it from proteasomal degradation. Importantly, the stabilised HIF-α, while still prolyl-hydroxylated, is transcriptionally active. We also showed that even the oxygen-insensitive form of HIF-α, HIF-α DM^(PP/AA)^, could still be degraded by the proteasome upon poly-ubiquitination. Furthermore, USP29 is able to reverse this ubiquitination and extend the half-life of the protein. The biological significance of this deubiquitination event is exemplified by the finding that USP29 expression levels correlate with the expression of the HIF target gene *CA9*, as well as with disease progression and severity in prostate cancer samples.

Most studies on HIF signalling are focused on HIF-1α and little is known about DUBs altering HIF-2α expression in spite of the functional divergence of both isoforms (Gonzalez-Flores et al, 2014). So far there is only one report showing that Cezanne/OTUD7B indirectly regulates *EPAS1* transcript through the regulation of E2F1 expression but there is no information about DUBs regulating HIF-2α stability (Moniz et al, 2015). Here we show for the first time that USP29 exhibited ubiquitin hydrolase activity towards HIF-1α and also HIF-2α in a similar way.

While a few UPS related negative and positive non-canonical regulators of HIF-α stability were described in the last few years (Schober & Berra, 2016), none of them has been shown to target HIF-α on its C-terminal end where we found USP29 to act on. We have identified a cluster of three lysine residues (K752, K755 and K758) located at the very C-terminal tail of HIF-1α as potential USP29 target site(s). As a matter of fact, the mutation of these residues to arginine almost completely abolished the basal ubiquitination and stabilised the mutated protein. However, we were unable to confirm by mass spectrometry that any of those lysines were indeed ubiquitinated as they weren’t resolved in the analysis, even though K48-linked polyubiquitin was present in the samples. The relevant sequence context suggested that fragmented peptides were either too long or too short to be resolved by MS.

Our attempts to identify the ubiquitin E3 ligase that ubiquitinates HIF-α on those lysines and therefore counteracts USP29 function have so far been unsuccessful. Nevertheless, we hypothesise that this Ub E3 ligase might have a crucial role in triggering HIF-α proteasomal degradation in a prolyl-hydroxylation-independent manner and could switch HIF signalling off even in hypoxic conditions. The phosphorylation of HIF-1α by ATM and PKA at S692 and S696, respectively has been suggested to increase its stability (Bullen et al, 2016; Cam et al,2010). Even though the effect of these kinases on HIF-2α has not been reported, the close proximity of the serine residues to the USP29-targeted lysines make it tempting to speculate that phosphorylation might increase or decrease the binding of USP29 and the relative Ub E3 ligase to HIF-α, respectively, and thereby determine HIF-α’s ubiquitination pattern and consequent stability.

To date, USP29 has been reported to exhibit deubiquitinase activity towards p53 and claspin, both proteins that are associated with carcinogenesis. The novel effect we now report on HIF-α protein levels expands the impact of USP29 in cancer. Since USP29 is involved in the regulation of key cellular processes such as HIF signalling and DNA integrity, it is not surprising to find USP29 expression to be very tightly controlled in healthy cells, transcriptionally and also potentially post-translationally. USP29, also known as HOM-TEST-84/86, is an imprinted gene located on chromosome 19q13.43 and encodes a protein of 922 Aa (Kim et al., 2007). As its neighbouring gene *Peg3*, USP29’s maternal allele is inactivated by imprinting and as a consequence, we and others found endogenous USP29 mRNA and protein levels barely above background by qPCR and Western Blot, respectively, using commercially available antibodies for protein detection (Kim et al, 2000; Liu et al, 2011). However, the silencing of endogenous USP29 by RNAi clearly affected HIF-α ubiquitination, suggesting that although being scarce, USP29 was catalytically highly active. The epigenetic mechanisms that control USP29 expression and how those mechanisms are disturbed in cancer remain to be determined. For instance, LOI (loss of imprinting)-mediated activation of the normally silent maternal allele might cause the USP29 upregulation, which we found in prostate cancer relative to non-tumour tissues. Interestingly, Liu and co-workers suggested that *USP29* expression was induced upon oxidative stress (Liu et al, 2011). In their experimental setup H_2_O_2_ treatment induced cooperative binding of FBP (FUSE binding protein) and AIMP2 (JTV1/p38) to *USP29*’s Far Upstream Sequence Element (FUSE), thereby triggering *USP29* transcription. Notably, AIMP2-DX2, an AIMP2 splice-variant, was particularly effective in inducing *USP29* expression (Liu et al, 2011) and high AIMP2-DX2 expression has been correlated with lung cancer progression (Choi et al, 2011).

The identification of USP29 as an important regulator of HIF-α provides a novel mechanism to explain the constitutive expression of HIF-α reported in many tumours independently of oxygen availability. Overexpression or hyperactivity of USP29 would therefore cause sustained HIF signalling, for which we found evidence in prostate cancer. However, it remains to be confirmed whether in these tumours HIF deubiquitination is indeed abnormally regulated. Alternatively, the loss of the respective so far unknown Ub E3 ligase or the mutation of the USP29 target lysines could provide a selective advantage for tumour cells. In this regard, a thorough sequencing effort in a broad range of tumours is needed to determine whether the site that we identified is mutated in human cancers and whether mutations evolve in metastatic progression or with drug-resistance. We found the mRNA expression levels of USP29 being associated with GS in prostate cancer, which suggests that USP29 may potentially serve as a prognostic marker. Finally, our results suggest that USP29 inhibitors could be used to switch HIF signalling off as a useful strategy in combination with current chemotherapies. In this context, it is worthy to note that USP29, having a cysteine protease catalytical site, is a potentially druggable protein.

Taken together, our study provides a rationale to make USP29 an important target for future studies. The further characterisation of the enzyme, its regulation and target proteins are crucial steps in order to understand how to tackle its deregulation.

## Materials and methods

### Plasmids

HIF-1α^630-826^ was amplified via PCR from pCMV-Myc-HIF-1α and inserted into the BamHI/ApaI-digested pCMV-Myc-HIF-1α vector. Green fluorescent protein-tagged HIF-1α DM^(P402/564A)^ was generated by inserting the sequence of Clover (Addgene plasmid #40259) behind the myc-tag in the BamHI-digested pCMV-Myc-HIF-1α DM^(PP/AA)^ construct (Berra et al,2003) using In-Fusion HD Cloning (Clontech). Then, replacing HIF-1α DM^(PP/AA)^ with HIF-2α DM^(PP/AA)^ generated green fluorescent HIF-2α DM^(PP/AA)^. mCherry-USP29 was generated by inserting the PCR-amplified mCherry sequence (Shaner et al, 2004) between the BspEI and NheI restriction sites of GFP-USP29. HIF-α truncations (HIF-1α^630-713^ and HIF-1α^630-750^) as well as HIF-1α DM^K752/755/758R^, HIF-2α DM^(P405/531A)^, HA-USP29 siRNA-resistant and the catalytically inactive USP29^C294S^ were generated using the QuikChange^®^ II XL Site-Directed Mutagenesis Kit (Stratagene) and the oligos reported in Table 1. All of the constructs were verified by sequencing. CMV-β-galactosidase and pRE-Δtk-Luc-HRE have been described before (Berra et al, 2003) as well as HIF-2α-Myc (Tian et al, 1997). Pools of shRNAs are from Open Biosystems (pSM2c-shRNA library (Silva et al, 2005)). The FLAG-ubiquitin plasmid was a gift from (Lee et al, 2014). HA-USP29 and GFP-USP29 expression vectors were gifts from (Liu et al, 2011)

**Table 1:**
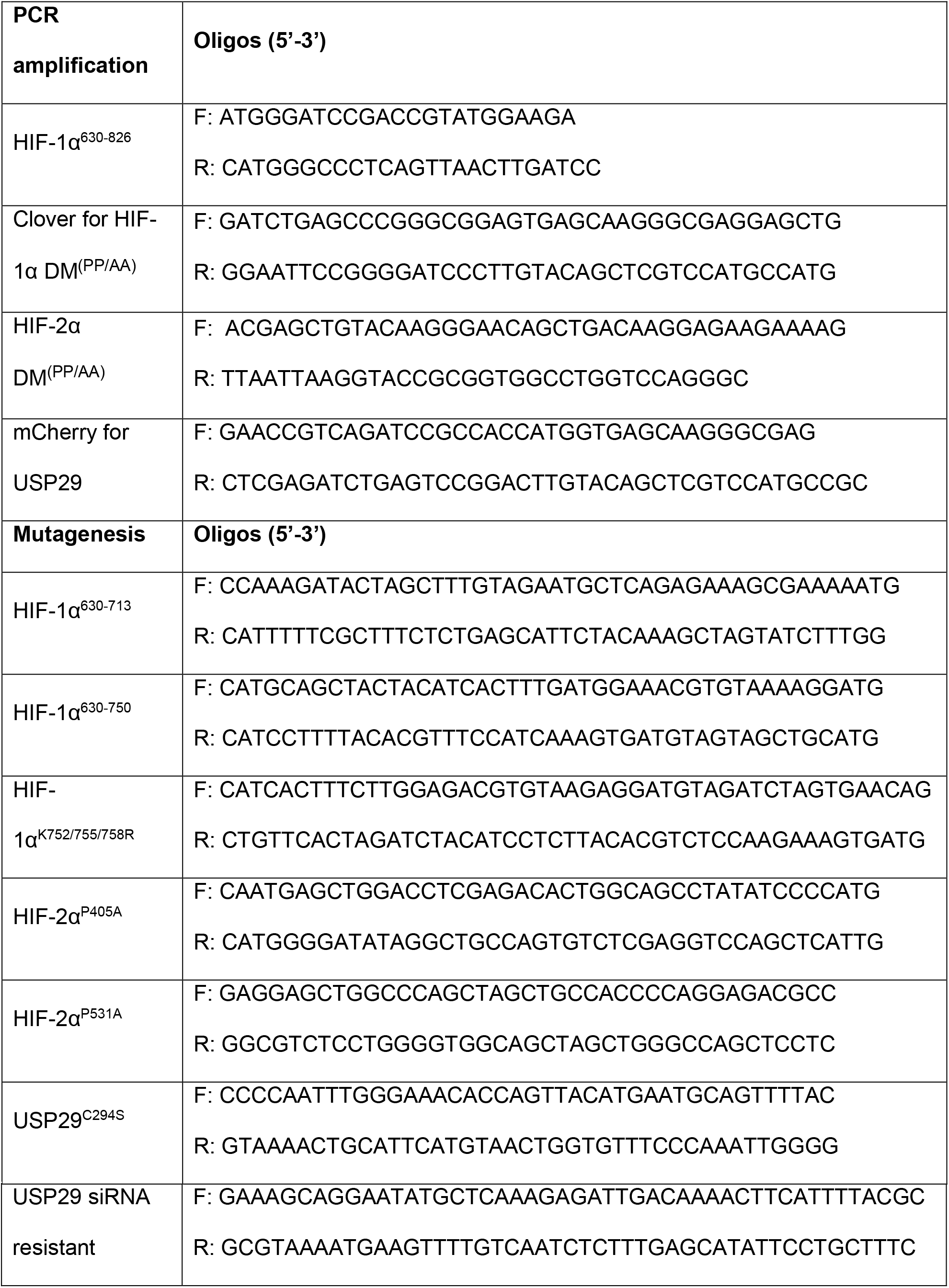
Oligo sequences for cloning and mutagenesis of HIF-α and USP29 plasmids.

### Cell culture and transfections

HEK293T cells were cultured in Dulbecco’s modified Eagle medium (DMEM) supplemented with 5% FBS. HeLa and PC3 cells were cultured in DMEM supplemented with 10% FBS and SK-N-AS cells with 1% non-essential amino acids additionally. A2780 and LNCaP were cultured in RPMI supplemented with 10 % FBS, and MDA-MB-231 and SH-SY5Y cells were cultured in DMEM:F12 (1:1) supplemented with 10% FBS. Cells were incubated at 37°C at 95% humidity and 5% CO_2_.

For delivery of siRNA or DNA to the cell, cells were transfected in suspension at plating or 24 h post-seeding at 60–70% confluence, respectively, using Lipofectamine 2000 (Invitrogen) as a transfection reagent following manufacturer’s instructions (Table 2 summarizes the sh- and siRNA sequences used in the manuscript). Incubation in hypoxia was achieved in an anaerobic workstation (*In vivo*_2_ 400, Ruskinn) and cell lysis was performed inside the anaerobic workstation to avoid reoxygenation.

**Table 2:**
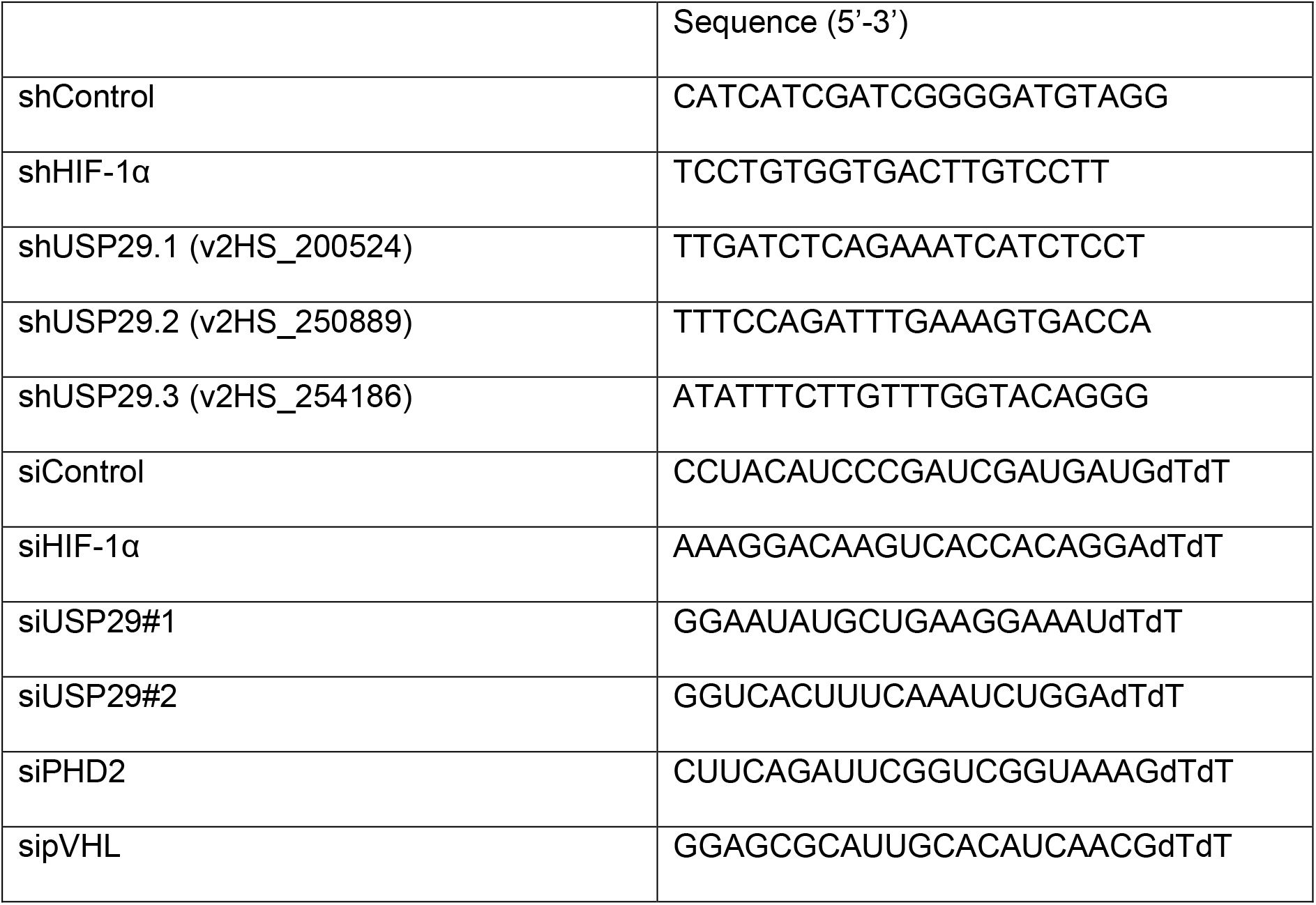
Sequences of shRNAs and siRNAs.

### Reporter assays and qRT-PCR

Cells were lysed in 25 mM Tris phosphate pH 7.8, 8 mM MgCl_2_, 0.5% Triton X-100, 7.5% glycerol and 1 mM DTT. Luciferase activity measurement was performed using the Steadylite plus™ High Sensitivity Luminescence Reporter Gene Assay System (PerkinElmer). β-galactosidase activity measurement was performed using the Galacto-Light Plus system (Applied Biosystems).

Total RNA was isolated using the RNeasy Mini Kit (Qiagen), reverse transcribed with qScript cDNA SuperMix (Quanta Biosciences) and primer-specific amplified with the quantitative PCR MasterMix FastStart Universal SYBR Green (Roche) or the TaqMan^®^ Universal Master Mix II when using the Universal Probe Library (Roche). The primer sequences and probes are listed in Table 3. PCR was carried out in a CFX96™ Thermal cycler (Bio-Rad). The expression of each target mRNA relative to *RPLP0* was calculated based on the threshold cycle (Ct) as 2^-Δ(ΔCt)^.

**Table 3:**
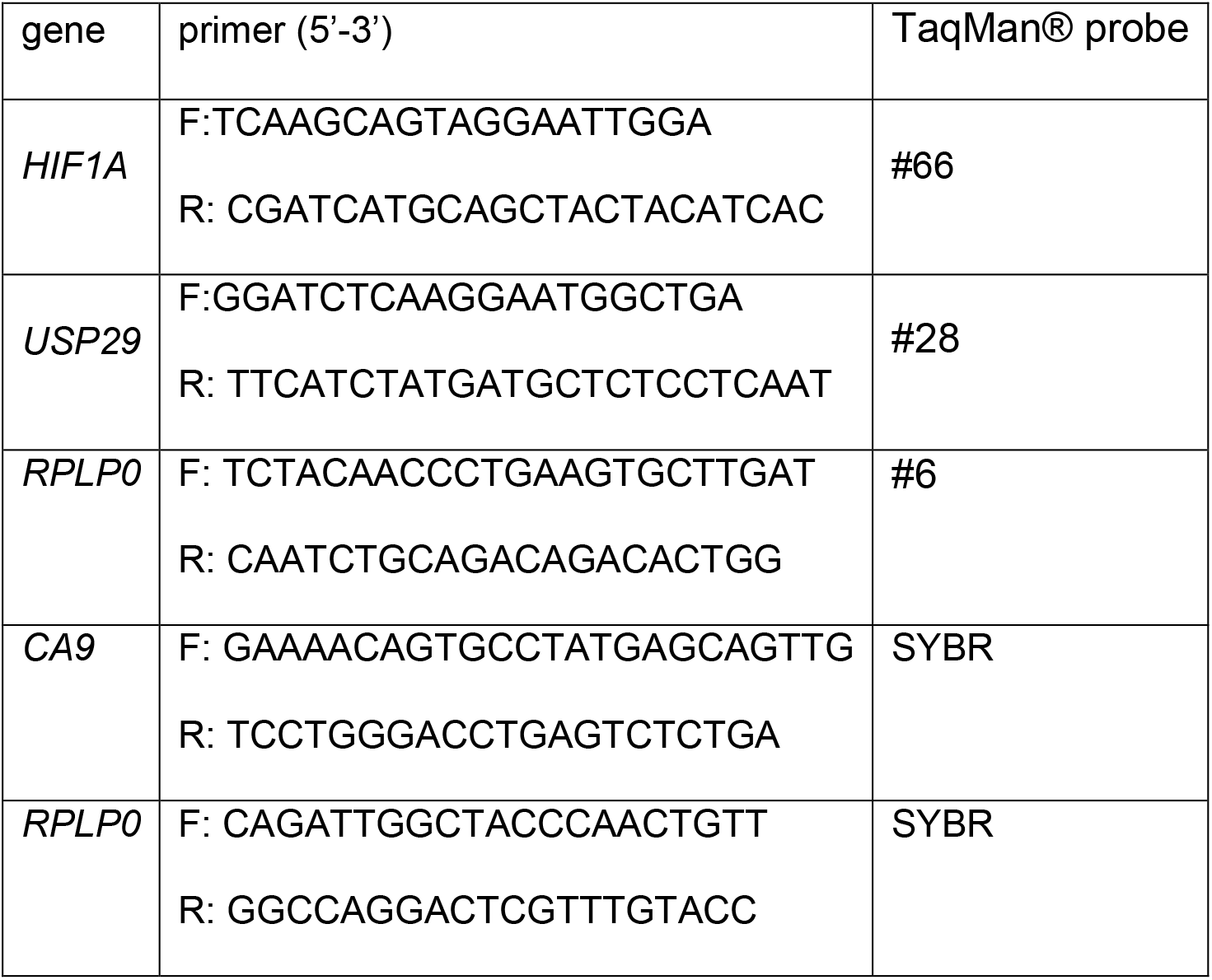
Sequences of primers and probes for qPCR.

### Ubiquitination assay, co-immunoprecipitation and immunoblotting

Ubiquitination assays were performed as described previously (Lee et al, 2014). Essentially, HEK293T cells were co-transfected with FLAG-tagged ubiquitin together with the expression vector of the GFP-tagged protein of interest. Cell were treated with MG132 (10 μM) for 2h prior to lysis with lysis buffer (50 mM Tris-HCl pH 7.5, 150 mM NaCl, 1 mM EDTA, 0.5% Triton X100, 40 mM β-Glycerolphosphate, 1 μg/ml Leupeptin, 1 μg/ml Aprotinin, 1 μg/ml Pepstatin A, 7 mg/ml N-ethylmaleimide (NEM)). Precleared lysates were incubated for 2.5h at RT with pre-washed GFP-traps^®^ (Chromotek) and subsequently subjected to stringent washes in denaturing conditions (8 M urea, 1% SDS). Protein was eluted by boiling at 95°C for 5 min (250 mM Tris-HCl pH 7.5, 40% glycerol, 4% SDS, 0.2% bromophenol blue, 5%β-mercaptoethanol) and migrated on 4-15% Tris-glycine gradient gels (BioRad).

Co-immunoprecipitation assays were performed as ubiquitination assays but cells were lysed in the absence of NEM (50 mM Tris-HCl (pH 8), 120 mM NaCl, 1 mM EDTA, 1 % IGEPAL CA-630, 40 mM β-Glycerolphosphate, 1 μg/ml Leupeptin, 1 μg/ml Aprotinin, 1 μg/ml Pepstatin A). Lysates were precleared by incubating with agarose beads (Chromotek) prior to overnight incubation with the GFP-traps^®^ and mild washes were performed with detergent-free non-denaturing lysis buffer. Protein complexes were eluted and migrated as described above.

For total cell extracts, cells were lysed with Laemmli buffer (50 mM Tris-HCl pH 6.8, 1.25% SDS, 15% glycerol) and total protein was quantified using the Lowry assay. Proteins were separated by SDS-PAGE and transferred onto a PVDF membrane (Millipore). The following antibodies were used for immunoblotting: mouse anti-β-actin (A5441, Sigma-Aldrich), mouse anti-CAIX (clone MN75, Bayer), mouse anti-FLAG-HRP (F3165, Sigma-Aldrich), mouse anti-GFP (11814460001, Roche), mouse anti-HA.11 (MMS-101R, Covance), rabbit anti-HIF-P564OH (D43B5, Cell Signaling Technology), anti-LC3 (2775s, Cell Signaling Technology), mouse anti-myc (9B11, Cell Signalling Technology). Home made rabbit anti-HIF1α and anti-PHD2 antibodies have been previously described (Berra et al, 2003). Immunoreactive bands were visualized with ECL.

### FLIM-FRET

Fluorescence lifetime images were acquired by scanning the sample with the LSM780 (Zeiss) scan head unidirectional and without averaging, recording a frame of 256 x 255 pixel with a pixel dwell time of 25.21 μs. Excitation of the green-fluorescent donor fluorophore was controlled by the PDL 828 “Sepia II” unit (PicoQuant) operating the 485 nm pulsed diode laser (PicoQuant) with a repetition rate of 40 MHz. The objective used was a C-Apochromat 40x/1.2 W Corr M27 (Zeiss). Fluorescence emission was collected through a 520/535 nm bandpass filter onto the Hybrid Detector PMA 40 (PicoQuant). Exact time between laser excitation and photon arrival was recorded by the Time-correlated single photon counting device (TCSPC) TimeHarp260 (PicoQuant) and plotted in a histogram, thereby building up a fluorescence decay curve. An instrument response function (IRF) using erythrosine B was recorded in the same measurement conditions on an everyday basis as described previously (Szabelski et al,2009). SymPhoTime 64 software (PicoQuant) controlled all PicoQuant hardware devices and was used for analysis. All photons within the region of interest were included in lifetime fitting analysis. The TCSPC-curve was reconvoluted with the IRF and fitted to a two-component decay curve to extract average lifetimes T_Av Int_.

### Bioinformatics analysis and statistics

Bioinformatic patients analyses were performed as reported (Torrano et al. 2016). Data was retrieved from (Taylor et al, 2010). These data have been subjected to background correction, log2 transformation and quartile normalisation. For the comparison of gene expression levels between different pathophysiological status, normal distribution and variances were analysed and a parametric ANOVA test was applied. For correlations analysis, a Pearson correlation test was applied. Pearson’s coefficient (R) indicates the existing linear correlation (dependence) between two variables X and Y, giving a value between +1 and −1 (both included), where 1 is total positive correlation, 0 is no correlation, and −1 is total negative correlation. The p-value indicates the significance of this R coefficient. The confidence level used in this case was also of 95% (alpha value = 0.05).

A minimum number of three independent experiments were performed. Data represent mean ± s.e.m. of pooled experiments with the exception of the western blots that correspond to a representative replicate. For data mining analysis, ANOVA test was used for multi-component comparisons and Student *T* test or Mann Whitney *U* test for two-group parametric or non-parametric comparisons, respectively. The confidence level used for all the statistical analyses was of 0.95 (alpha value = 0.05).

## Acknowledgements

We would like to thank Dr Russell and Dr Levens for kindly providing the original HIF-2α and USP29 plasmids, respectively. EB’s lab is supported by the Basque Department of Industry, Tourism and Trade (Etortek) and Education (PI2010/16), and the MINECO (BFU2013-46647-R and BFU2016-76872-R) grants. ASS is a recipient of a UoL-CIC bioGUNE PhD studentship. TMM is supported by the MINECO FPI fellowship (BES-2014-070406). An ERC Starting grant (336343) supports AC’s lab and UM is a recipient of a MINECO grant (SAF2013-44782-P). We thank the Centre for Cell Imaging (CCI) for technical support. CCI equipment was funded by the MRC (MR/K015931/1). The authors declare that they have no conflict of interests.

## Author contributions

ASS designed and performed experiments and contributed to the analysis of the data and the writing of the manuscript. IMB, TMM, EPA, OC and SP performed experiments. ARC, AMA and AC carried out the bioinformatic and biostatistical analysis and contributed to the discussion of the results. UM and VS provided technical support and contributed to the critical discussion of the results and manuscript revision. EB designed and supervised the project, analysed data and wrote the manuscript.

## Conflicts of interest

The authors declare that they have no conflict of interests.

