## Supplemental data for "USP29 is a novel non-canonical Hypoxia Inducible Factor-α activator"

### Supplemental Figure legends

**Supplemental Figure 1: USP29 is a positive regulator of HIF-1α.**

**A** HeLa cells were silenced with scrambled or shRNAs targeting *USP29* and transfected with GFP-USP29. RNA and protein were extracted and subjected to qPCR and immunoblotting, respectively.

**B** HeLa cells were silenced with scrambled shRNAs or shRNAs targeting *HIF1A* and *USP29* and incubated for 16 h in normoxia (21% $O_2$) or hypoxia (1% $O_2$). Total RNA was extracted, reverse-transcribed and expression of *HIF1A* was determined by qPCR.

**C** HEK293T cells were transfected with empty vector or HA-USP29, RNA was extracted and expression of *USP29* as well as *HIF1A* was determined by qPCR.

**Supplemental Figure 2: USP29 regulates HIF-1α in a non-canonical way.**

**A** HEK293T cells were silenced with scrambled siRNA or two independent siRNA sequences (20nM) targeting *USP29* and transfected with myc-HIF-1α $DM^{(PP/AA)}$ or HA-USP29. WCE were analysed by WB for the expression levels of myc-HIF-1α $DM^{(PP/AA)}$ or HA-USP29, respectively.

**B** HEK293T cells were co-transfected with myc-HIF-1α or myc-HIF-1α $DM^{(PP/AA)}$ and HA-pVHL. Total cell extracts were subjected SDS-PAGE followed by immunoblotting with the indicated antibodies.

**Supplemental Figure 3: USP29 stabilises HIF-1α by protecting from proteasome-mediated degradation.**

**A** HEK293T cells were transfected with myc-HIF-1α DM$^{(PP/AA)}$ and left untreated or treated with either the proteasome inhibitor MG132 (10 μM), the autophagy inhibitor be 3 chloroquine (30 μg/mL) or both inhibitors together for 6 hours. Protein levels were determined by immunoblotting of the whole cell extracts with the indicated antibodies.

**B** HEK293T cells were co-transfected with myc-HIF-1α DM$^{(PP/AA)}$ and empty vector or GFP-USP29 and treated with cycloheximide (CHX) (20 μg/ml) to inhibit protein synthesis. Cell extracts were collected at the indicated times after CHX addition and subjected to immunoblotting with the indicated antibodies.

**Supplemental Figure 4: USP29 deubiquitinates HIF-α DM$^{(PP/AA)}$.**

**A** HeLa cells were transfected with the FRET donor HIF-2α DM$^{(PP/AA)}$-GFP alone or together with the FRET acceptor mCherry-USP29. Fluorescence images for donor (green) and acceptor (red) channel were acquired (left and central panel). The lifetime of the donor was measured and pseudo-colour coded fluorescence life time images (FLIM) were generated. Average lifetimes of the donor in the absence (n = 25) and the presence (n = 29) of the FRET acceptor were calculated from 3 independent experiments. Scale bars are 10 μm long, (*) $p = 2.36*10^{-11}$.

**B** HEK293T cells were co-transfected with HA-USP29 and either GFP alone or GFP-HIF-1α DM$^{(PP/AA)}$. Cells were lysed in native conditions and GFP-tagged protein was immunoprecipitated with GFP-traps®. Immuno-complexes were analysed for the presence of HA-USP29 by immunoblotting.

**C** HEK293T cells were co-transfected with GFP-HIF-2α DM$^{(PP/AA)}$, FLAG-ubiquitin and either HA-USP29 or empty vector. Cells were treated with the proteasome inhibitor MG132 (10 μM) for 2 hours and lysed in the presence of the DUB inhibitor NEM. GFP-

HIF-2α DM$^{(PP/AA)}$ was pulled down with GFP-traps® and subjected to stringent washes (8 M urea, 1% SDS). Ubiquitinated and non-ubiquitinated GFP-HIF-2α DM$^{(PP/AA)}$ protein in the eluate was analysed by immunoblotting with anti-FLAG and anti-GFP antibodies, respectively.

**Supplemental Figure 5: USP29 targets the C-terminal part of HIF-α.**

**A** HEK293T cells were co-transfected with myc-HIF-1α DM$^{630\text{-}826}$ and either empty vector, GFP-USP29 or GFP-USP29$^{C/S}$. Whole cell extracts were prepared and submitted to immunoblotting with the indicated antibodies.

**B, C** HEK293T cells were co-transfected with (B) myc-HIF-2α DM$^{601\text{-}870}$ or (C) myc-HIF-1α DM$^{630\text{-}826}$, myc-HIF-1α DM$^{630\text{-}713}$ or myc-HIF-1α DM$^{630\text{-}750}$ and either empty vector or GFP-USP29. Whole cell extracts were prepared and submitted to immunoblotting with the indicated antibodies.

**D** Alignment of the lysine-containing C-terminal sequence of HIF-1α and HIF-2α from human (H), mouse (M), rat (R), cow (T), xenopus (X) and zebrafish (Z).

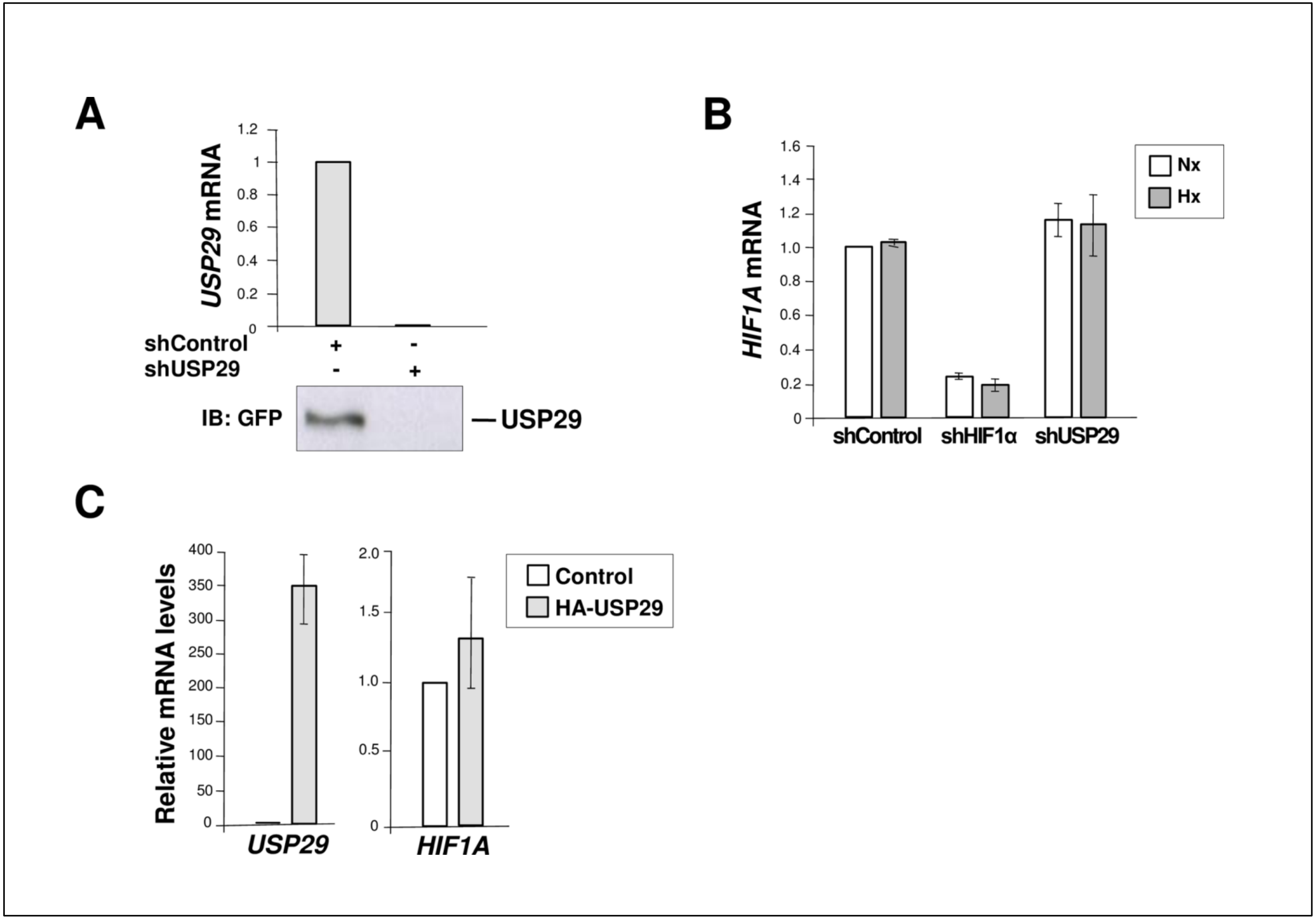


**Supplemental Figure 1**

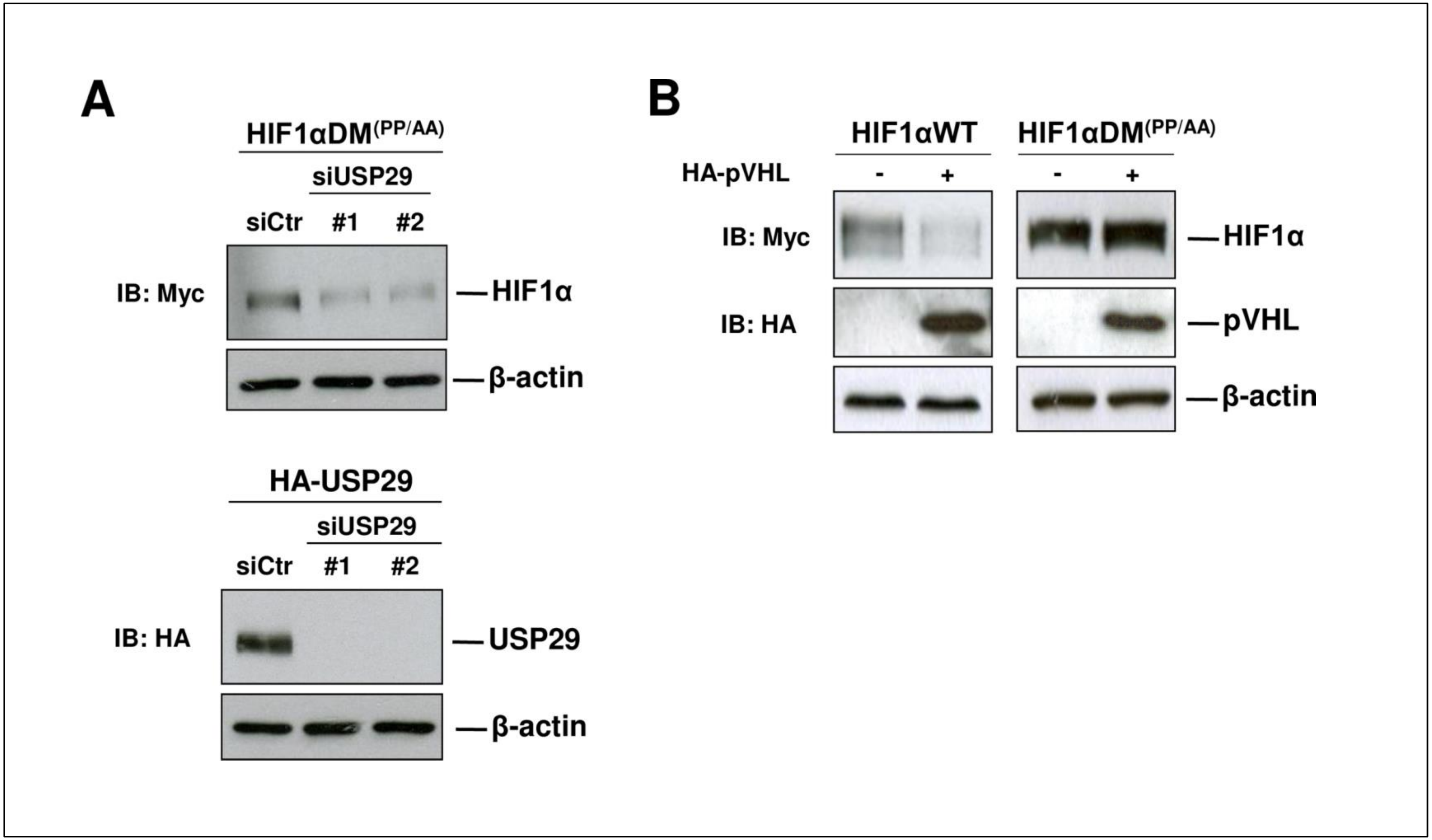


Supplemental Figure 2

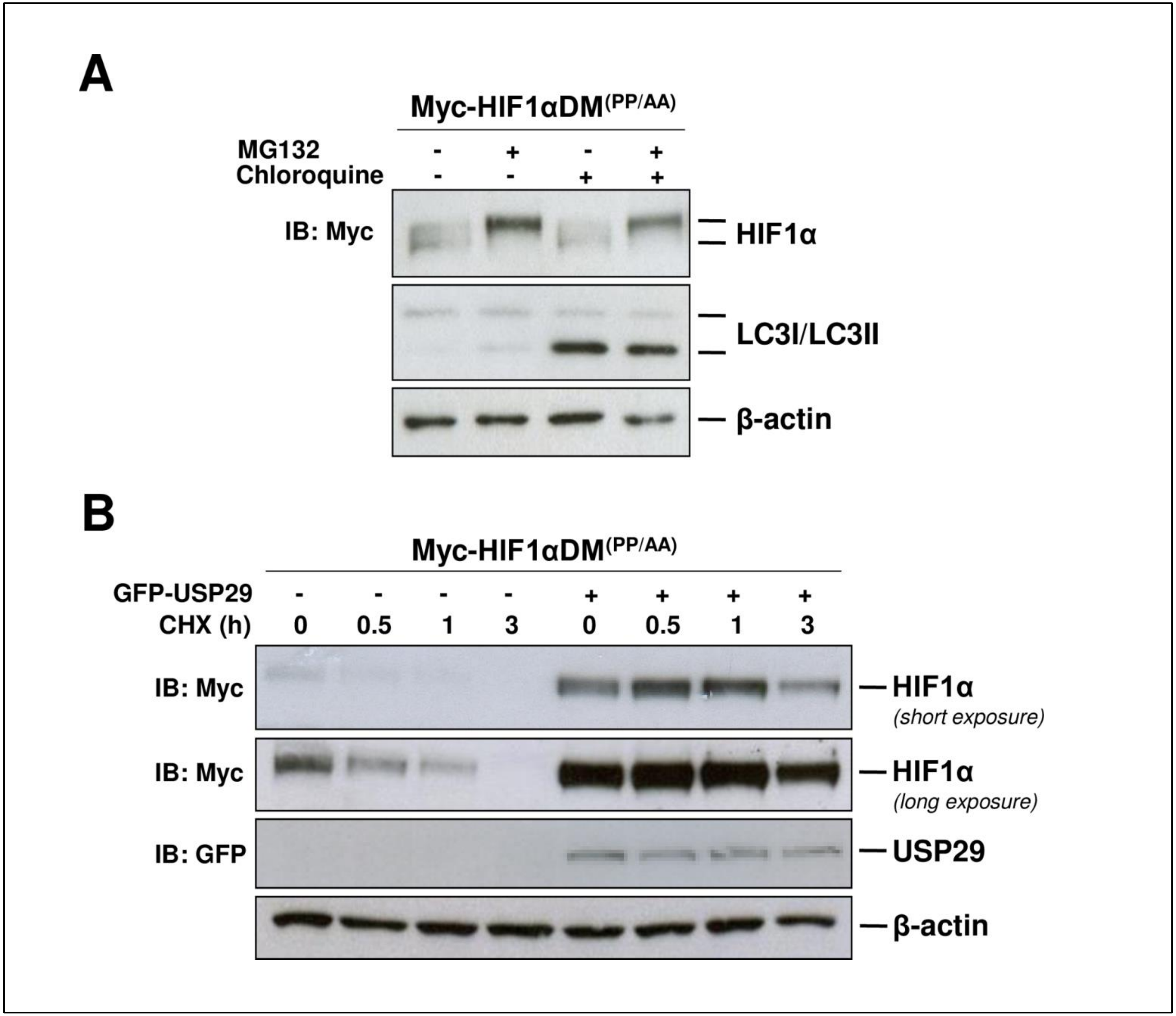


Supplemental Figure 3

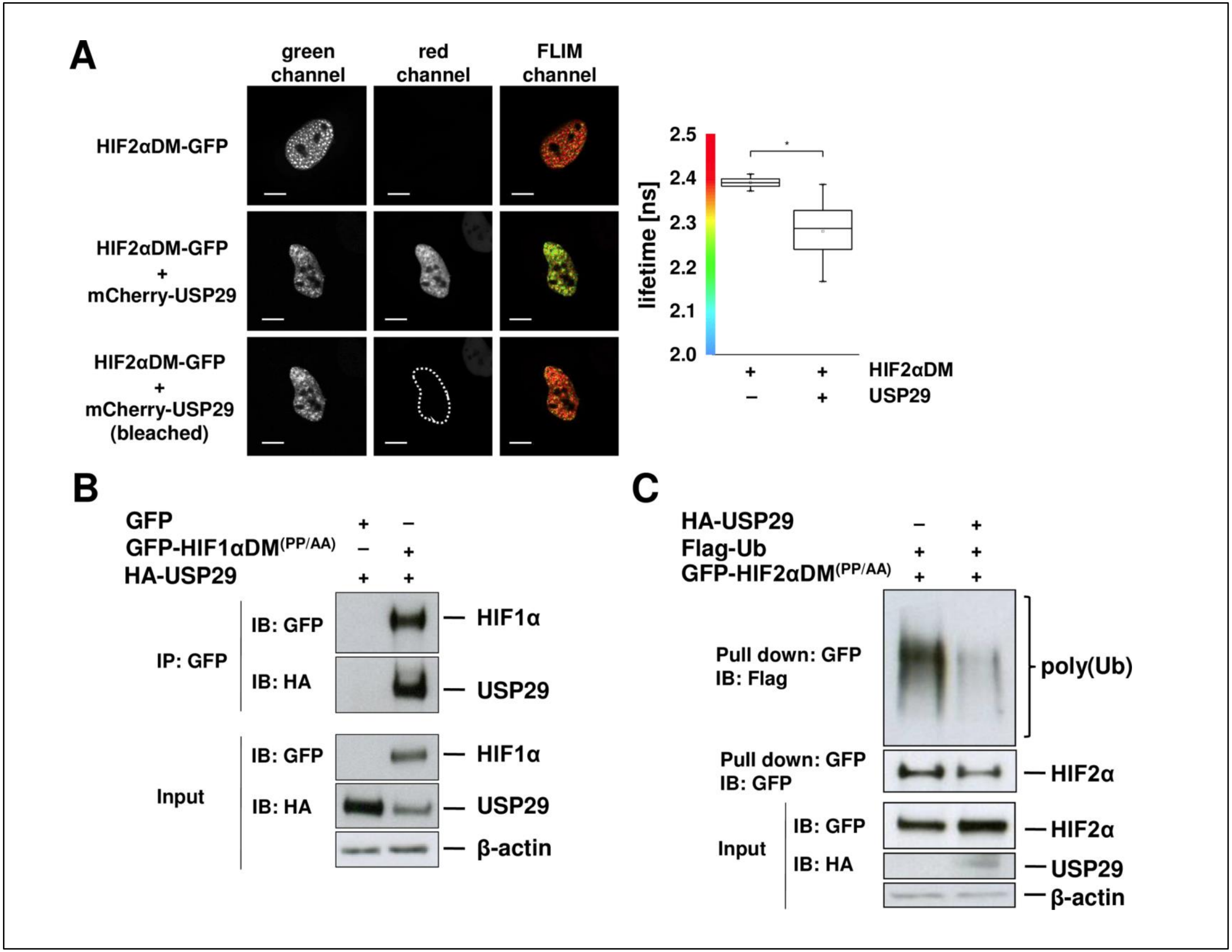


Supplemental Figure 4

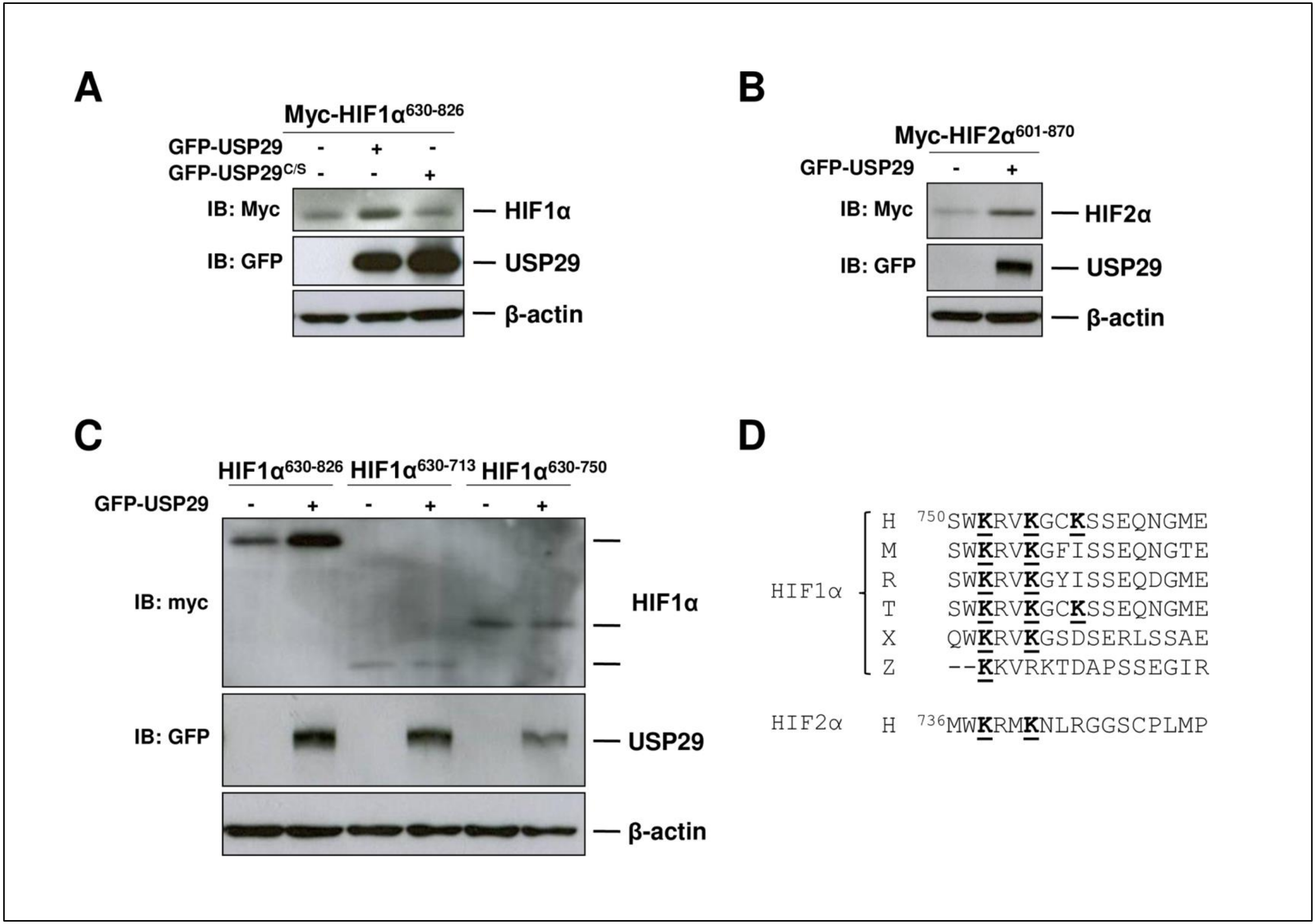


**Supplemental Figure 5**
